# Identification of the pathway of Rhodoquinone biosynthesis in *C.elegans*

**DOI:** 10.1101/627737

**Authors:** Samantha Del Borrello, Margot Lautens, Kathleen Dolan, June H. Tan, Mark A. Spensley, Amy A. Caudy, Andrew G. Fraser

**Author notes:** = these authors contributed equally (Samantha Del Borrello, Margot Lautens).

## Abstract

Parasitic helminths infect over a billion humans. To survive in the low oxygen environment of their hosts, these parasites use unusual anaerobic metabolism. This requires Rhodoquinone (RQ), an electron carrier that is made by very few animal species — crucially it is not present in any parasitic hosts. RQ synthesis is thus an ideal target for anthelmintics but little is known about how RQ is made and no drugs are known to block RQ synthesis. *C.elegans* makes RQ and can use RQ-dependent metabolic pathways — here, we use *C.elegans* genetics to identify the pathway for RQ synthesis and show that *C.elegans* requires RQ for survival in hypoxic conditions. Finally, we establish a robust assay for drugs that block RQ-dependent metabolism. This study identifies for the first time how RQ is made in any animal and establishes a novel assay that can drive the development of a new class of anthelmintic drugs.

## Introduction

Soil-transmitted Helminths (STHs) are major human pathogens (World Health Organization, 2002). Over a billion humans are infected with an STH — the roundworm *Ascaris lumbricoides*, the whipworm *Trichuris trichuria*, and the hookworm *Necator americanus* account for most of these infections (World Health Organization, 2002). STHs are transmitted from human to human via the soil where eggs from human faeces develop into infective stages which then enter new hosts (reviewed in Brooker et al., 2006). On infection, STHs encounter a very different environment and require multiple strategies to be able to survive. One of the major changes is the availability of oxygen — while there is abundant oxygen outside their hosts, in many host tissues there is little available oxygen and the parasites must switch from aerobic respiration to anaerobic respiration. Crucially, the anaerobic metabolic pathways that STHs depend on while in their hosts are unusual and are not used in any host (Klockiewicz et al., 2002). Inhibiting these anaerobic pathways thus provides a way to attack the parasites while leaving the host unaffected.

During aerobic respiration in helminths, the great majority of ATP is made in the mitochondrion (Tielens, 1994; Tielens et al., 1984). Electrons enter the Electron Transport Chain (ETC) either at Complex I or via several quinone-coupled dehydrogenases (QDHs from here on). These QDHs include Succinate Dehydrogenase (Complex II) and Electron-Transferring Flavoprotein Dehydrogenase (ETFDH) (Komuniecki et al., 1989; Ma et al., 1993; Rioux and Komuniecki, 1984). The electrons entering the ETC are first transferred to the lipid soluble electron carrier Ubiquinone (UQ) (Crane et al., 1957; Mitchell, 1975). From UQ, they are ultimately carried to Complex III then IV where they are finally transferred onto oxygen as the terminal electron acceptor (see Fig 1a). Electron transport is coupled to proton pumping into the inner membrane space of the mitochondrion — this establishes a proton gradient which is used to power the F0F1-ATP synthase (Mitchell, 1961). When there is insufficient oxygen to accept electrons at Complex IV, or when inhibitors of Complex IV such as cyanide (Antonini et al., 1971; Nicholls et al., 1972) are present, almost all animals stop using the ETC and rely on anaerobic glycolysis to make ATP, generating lactate (Isom et al., 1975; Meyerhof, 1927). STHs, however, have evolved a different solution that allow them to survive months in the hypoxic host environment.

**Figure 1:**
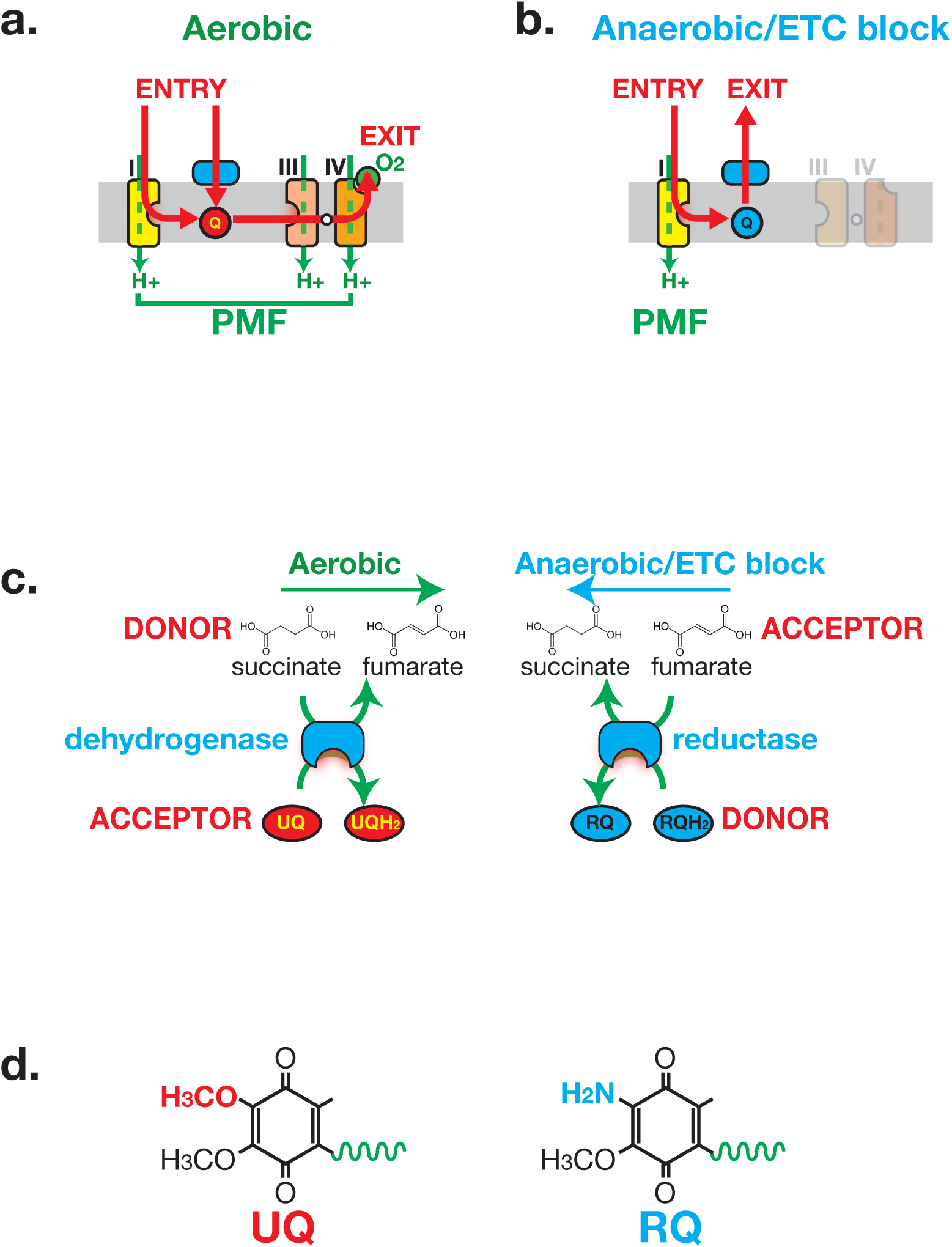
Anaerobic metabolism in helminths requires Rhodoquinone (RQ). **a. Electron flow in the Electron Transport Chain (ETC) under aerobic conditions.** Electrons enter the ETC either via Complex I or via a number of Quinone-coupled Dehydrogenases (QDH; cyan). These complexes transfer electrons to Ubiquinone (red circle ‘UQ’) which shuttles them to Complex III. They exit the ETC at Complex IV where they are transferred to oxygen as the terminal electron acceptor. Proton pumping is coupled to electron transport and is carried out by Complexes I, III and IV. Electron flow is shown in red and proton pumping in green. **b. Electron flow in the Electron Transport Chain (ETC) under anaerobic conditions.** Electrons still enter the ETC at Complex I which transfers electrons to RQ (cyan circle ‘RQ’). RQ shuttles electrons to the QDHs which now operate as reductases, allowing electrons to exit the ETC and onto a diverse set of terminal electron acceptors. Complex I is the sole proton pump in this truncated ETC. **c. Schematic of Complex II activity under aerobic and anaerobic conditions.** Under aerobic conditions, Complex II acts as a succinate dehydrogenase, transferring electrons from succinate onto UQ. Under anaerobic conditions, Complex II operates in the reverse direction acting as a fumarate reductase, accepting electrons from RQ and transferring them to succinate as the terminal electron sink. **d. Structure of UQ and RQ.** The critical amine group differing between UQ and RQ is highlighted; the prenyl tail is shown schematically as a green wavy line.

Electrons still enter the ETC at Complex I, Complex I still pumps protons to generate the proton motive force (PMF), and ATP is still made by the F0F1ATPase, powered by the PMF. However, rather than the electrons passing through the ETC to oxygen as the terminal electron acceptor, they exit the ETC immediately downstream of Complex I onto a number of alternative terminal electron acceptors (Fig 1b) (reviewed in Hochachka and Mustafa, 1972; Müller et al., 2012). This transfer of the electrons out of the ETC and onto alternative electron acceptors requires the quinone-coupled dehydrogenases (Kita, 1992; Ma et al., 1993). Under aerobic conditions these QDHs act as entry points to the ETC, transferring electrons from their substrates to UQ. Crucially, the reactions catalysed by these QDHs are reversed in anaerobic conditions — they now act as reductases transferring electrons out of the ETC and onto their products. For example, Complex II acts as a succinate dehydrogenase in aerobic conditions, generating fumarate; in anaerobic conditions, it reduces fumarate generating succinate as a terminal electron sink (Fig 1c) (Klockiewicz et al., 2002; Kmetec and Bueding, 1961; Sato et al., 1972; Saz and Vidrine, 1959; Takamiya et al., 2002). In this way an entry of electrons into the ETC from a variety of electron donors in aerobic conditions is reversed to provide an exit from the ETC onto a variety of electron acceptors in anaerobic conditions.

The unusual ETC wiring used by STHs to survive anaerobic conditions requires an unusual electron carrier, Rhodoquinone (RQ) (Moore and Folkers, 1965). RQ and UQ are highly related molecules — the sole difference is the presence of an amine group on the quinone ring of RQ (Fig 1d). This changes the biophysical properties of the quinone ring: while UQ can accept electrons from the QDHs as they flow into the ETC under aerobic conditions, UQ cannot carry electrons of the correct electropotential to drive the reverse reactions in anaerobic conditions (Fioravanti and Kim, 1988; Sato et al., 1972). RQ can carry such electrons (Fioravanti and Kim, 1988; Sato et al., 1972), however, and the ability of STHs to survive in their hosts is absolutely dependent on RQ. The single amine group that differs between UQ and RQ thus affects the health of over a billion humans. RQ is found in very few animal species — only nematodes, molluscs and annelids are known to make RQ (Allen, 1973; Fioravanti and Kim, 1988; Klockiewicz et al., 2002; Sato and Ozawa, 1969; Takamiya et al., 2002; Van Hellemond et al., 1995). Since no host animals make RQ, inhibiting RQ synthesis or RQ use is a potentially powerful way to target parasites inside their host. Currently however little is known about RQ synthesis. The most mature studies have focused on the purple Proteobacterium *R.rubrum*, where RQ appears to derives from UQ (Brajcich et al., 2010). RQ synthesis in *R.rubrum* requires the gene *rquA* (Lonjers et al., 2012) which is the first and thus far only gene known to be required for RQ synthesis in any organism. It is still unclear what role it plays in RQ synthesis (Lonjers et al., 2012), nor what the rest of the genes required for RQ synthesis may be. In animals, the situation is even more blank: nothing is known about which genes are required for RQ synthesis and there are no drugs that are known to prevent RQ synthesis. This is partly because no tractable animal model has been established in which to study RQ synthesis and use.

Previous studies have shown that *C.elegans*, a free-living helminth, can make RQ (Takamiya et al., 2002) and that when *C.elegans* is exposed to hypoxic conditions it undergoes major metabolic changes that resemble those that occur in STHs when they are in the hypoxic environment of their hosts (Butler et al., 2012; Föll et al., 1999). This suggested to us that we could establish *C.elegans* as a model for dissecting the pathway of RQ synthesis and for screening for drugs that block RQ synthesis or use. We confirm that *C.elegans* makes RQ and also that it uses RQ-dependent metabolism when unable to use oxygen as a terminal electron acceptor. Crucially, we identify the pathway of RQ synthesis in *C.elegans* and show that *C.elegans* requires RQ to survive under conditions where oxygen cannot be used as an electron acceptor for the ETC. This allowed us to establish a robust high throughput screening assay to identify compounds that block RQ synthesis or RQ use. This is the first study to show how RQ, an electron carrier that affects the life of over a billion humans, is made in helminths. This will help towards the development of a new class of drugs to treat these major human pathogens.

## Results

### *C.elegans* makes RQ and switches to anaerobic metabolism when exposed to Potassium Cyanide

*C.elegans* is a non-parasitic helminth that is easily genetically tractable (Brenner, 1974; reviewed in Jones et al., 2005) and can be used for efficient drug screens (Burns et al., 2015). We wanted to establish *C.elegans* as a model to dissect the pathway for RQ synthesis in helminths and as a system in which we could efficiently screen for drugs that block the synthesis of RQ or use of RQ. We note that there are no other standard model organisms where this is possible: yeasts, insects, fish, and vertebrates do not make or use RQ so *C.elegans* is the sole genetically tractable animal model for this work.

Like other helminths, *C.elegans* has previously been shown to make RQ (Takamiya et al., 2002). We wanted to confirm this and determine whether we could define a simple experimental method to drive *C.elegans* to carry out similar anaerobic metabolism as that used by parasitic helminths to survive in their hosts. We extracted quinones as described in Methods and as shown in Fig 2a and Supp Fig 1, *C.elegans* makes both UQ and RQ when maintained in normoxic conditions. We can therefore use *C.elegans* to genetically dissect the pathway for RQ synthesis.

**Figure 2:**
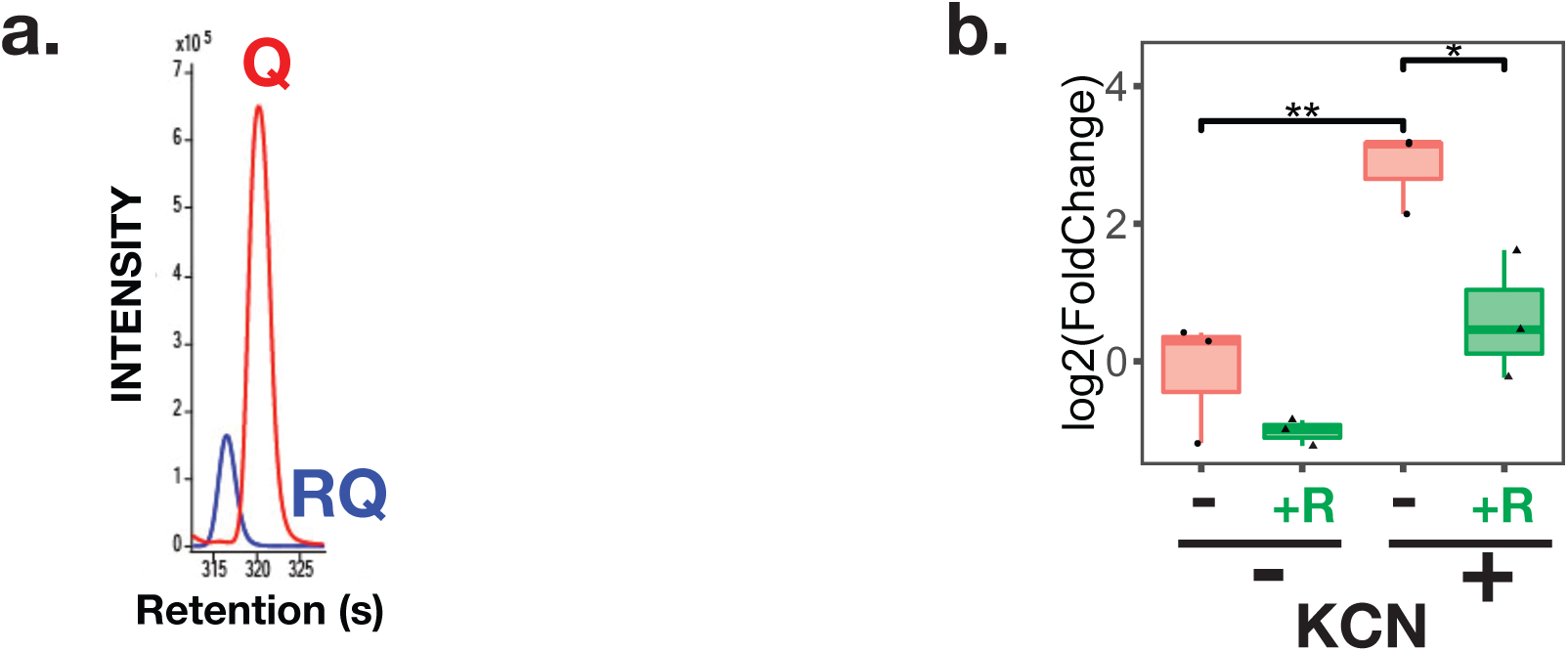
*C.elegans* makes Rhodoquinone (RQ) and can carry out RQ-dependent anaerobic metabolism. **a. *C.elegans* makes both UQ and RQ.** *C.elegans* were grown under normoxic conditions and quinones extracted and analysed by mass spectrometry (see Methods). Both UQ and RQ can be detected. **b. *C.elegans* increases succinate production following treatment with Potassium Cyanide (KCN).** *C.elegans* L1 larvae were treated either with 200 µM KCN alone, 12.5 µM rotenone alone, or a combination of KCN and rotenone for 1 hr and metabolites extracted and analysed by mass spectrometry (see Methods). The graph shows that succinate levels increase over 5-fold following KCN treatment and that this increase is blocked by rotenone indicating that it requires Complex I activity. Data are from 3 independent biological repeats; box plots show median and the interquartile range as well as individual datapoints.

Our next step was to establish a simple method to drive *C.elegans* to use RQ-dependent metabolism that would allow high throughput drug screens. Parasitic helminths use RQ-dependent metabolism under low oxygen conditions (Rioux and Komuniecki, 1984; Saz and Lescure, 1969; Tielens et al., 1992) and previous studies showed that *C.elegans* shows similar metabolic shifts when exposed to hypoxic conditions (Butler et al., 2012; Föll et al., 1999). If possible, however, we wanted to avoid the use of hypoxic chambers. While hypoxia chambers are highly accurate ways of controlling oxygen levels, they are also very expensive and cumbersome for large-scale drug screens. We therefore turned to chemical methods of inducing a hypoxic state. Potassium Cyanide (KCN) is a potent inhibitor of Complex IV (Antonini et al., 1971; Nicholls et al., 1972) — KCN inhibits oxygen binding to Complex IV and KCN treatment thus mimics hypoxia. We tested whether treatment with KCN could drive *C.elegans* to use anaerobic metabolism that is similar to the RQ-dependent metabolism used by STHs in their hosts. The classic hallmark of RQ-dependent anaerobic metabolism in helminths is the generation of high levels of succinate through the reversal of Complex II (Fig 1c) (Butler et al., 2012; Saz and Lescure, 1969; Tielens et al., 1992). If *C.elegans* can indeed use the same anaerobic metabolism as parasitic helminths, there should be a build-up of succinate following KCN treatment. Furthermore, this should be dependent on Complex I activity, since Complex I is the sole source of electrons that are carried by RQ to drive the fumarate reductase activity of Complex II (Fig 1b). We found that when *C.elegans* are exposed to KCN, they build up high levels of succinate as expected and that inhibiting Complex I with rotenone prevents succinate build-up (Fig 2b). We thus find that *C.elegans* makes RQ and that treatment of *C.elegans* with KCN causes them to switch to a metabolic state that resembles that of STHs in their host. *C.elegans* is thus an excellent model in which to dissect RQ synthesis and to screen for compounds that alter RQ-dependent metabolism.

### In *C.elegans*, RQ is synthesised from products of the kynurenine pathway and not from Ubiquinone

RQ and UQ are highly related molecules — the sole difference is the presence of an amine group on the quinone ring of RQ (Fig 1d). The critical question for RQ synthesis is where this amine group comes from and how it is generated. The best-defined current model for RQ synthesis comes from experiments in the proteobacterium *R.rubrum*. At least in this prokaryote, RQ is thought to be made by a late addition of the critical amine group to an existing molecule of UQ (Brajcich et al., 2010). UQ is thus an obligate precursor of RQ and RQ synthesis requires initial synthesis of UQ (Fig 3a). While this may be the case for *R.rubrum*, this is not the case in *F.hepatica* (Van Hellemond et al., 1996) or *C.elegans*. The *clk-1(qm30)* strain has a loss-of-function mutation in the *C.elegans* COQ7 orthologue that is required for hydroxylation of 5-demethoxyubiquinone to 5-hydroxyubiquinone, a late step in UQ synthesis — there is no detectable UQ in *clk-1(qm30)* homozygous animals. However, a previous study showed that there does appear to be RQ in this strain (Jonassen et al., 2001). If there is no UQ, but there is RQ, then RQ is not derived from UQ, at least in helminths. The two models for RQ synthesis thus differ fundamentally — in one UQ is an obligate precursor (Brajcich et al., 2010), in the other it is not (Jonassen et al., 2001). Since this is a fundamental result, we wanted to confirm this before trying to dissect the pathway of RQ synthesis. We thus extracted and analysed quinones from either wild-type worms or *clk-1(qm30)* homozygous animals. We find that while there is no detectable UQ in *clk-1(qm30)* mutants, there is abundant RQ and indeed we find that RQ levels are essentially unchanged (Fig 3b). We thus confirm that UQ is not an obligate precursor for RQ in *C.elegans*.

**Figure 3:**
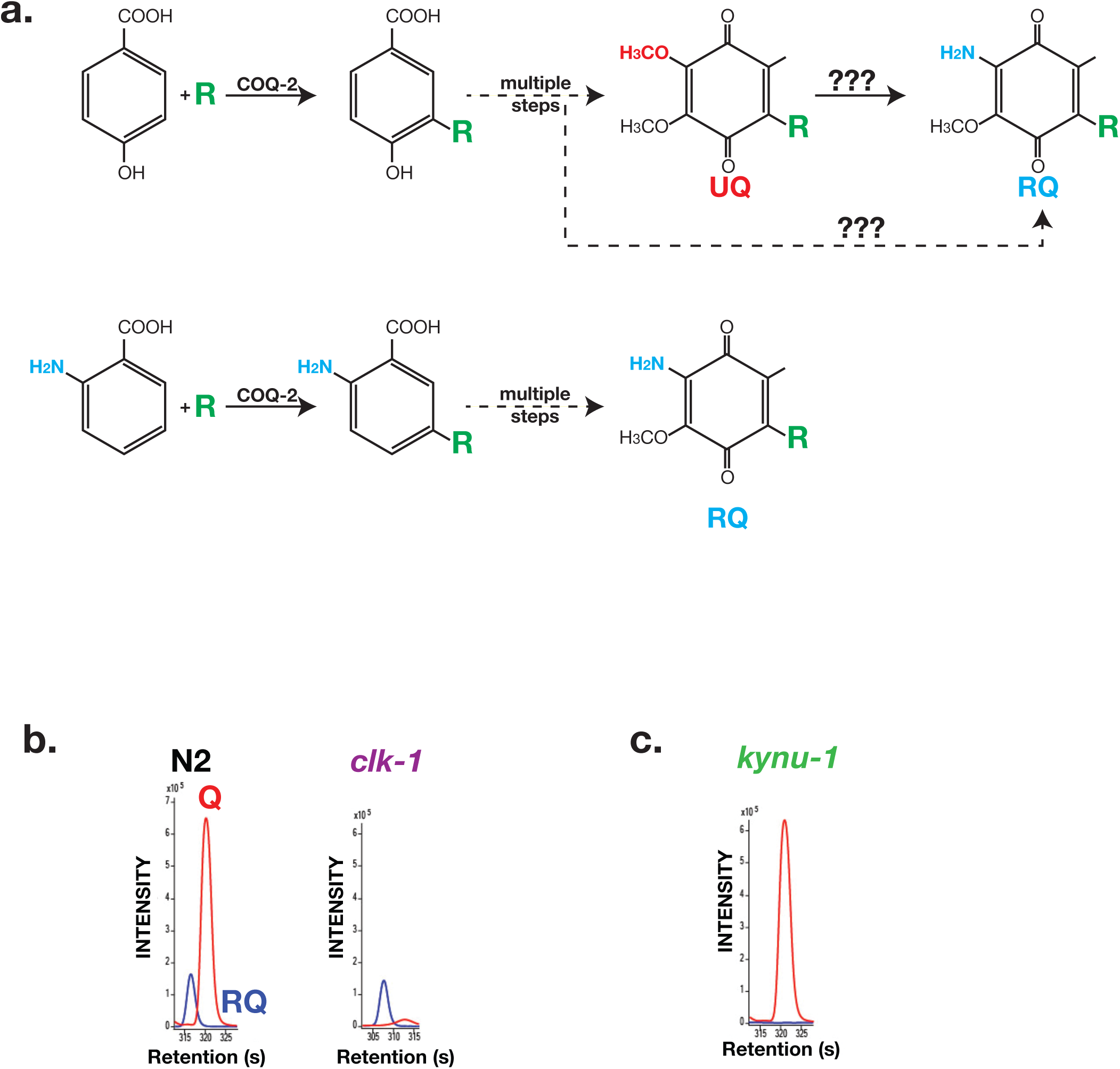
RQ in *C.elegans* does not derive from UQ but from Tryptophan metabolites. **a. Schematic showing possible routes for RQ synthesis.** Current models for RQ synthesis are shown schematically in the top pathway: PHB is prenylated by COQ-2 at the start of the UQ synthesis pathway and RQ either derives from UQ or from a UQ precursor — the amine group is thus added at a late step. The lower pathway shows our proposed pathway. Rather than use PHB as the substrate for COQ-2, RQ synthesis starts with prenylation of anthranilate or 3-hydroxyanthranilate by COQ-2 (the 3 hydroxy group is denoted by a red asterisk). The amine group is thus present from the start of RQ synthesis rather than being added at a late step and RQ synthesis proceeds via the same pathway as UQ synthesis. **b. UQ is not a required intermediate for RQ synthesis.** *clk-1(qm30)* mutant worms and N2 wild-type worms were grown under normoxic conditions and quinones extracted and analysed. N2 worms contain both UQ and RQ whereas *clk-1(qm30)* mutants only contain detectable RQ. **c. RQ synthesis requires metabolism of Tryptophan via the kynurenine pathway.** *kynu-1(e1003)* mutant worms were grown under normoxic conditions and quinones extracted and analysed. While wild-type worms contain both UQ and RQ, *kynu-1(e1003)* mutants only contain detectable UQ. RQ synthesis thus requires the generation of 3HA or anthranilate from Tryptophan via the kynurenine pathway.

If RQ is not generated by addition of the key amine group to an existing UQ molecule where does the amine group on RQ come from? One possibility is it is added not to UQ but to a UQ precursor such as demethoxyquinone (DMQ) — such UQ precursors would still be present in the *clk-1(qm30)* mutant strain (Fig 3a for schematic). While this could in principle be the case, it is unlikely because amination of an aromatic ring is highly thermodynamically unfavorable (reviewed in Downing et al., 1997). We therefore investigated an alternative possibility — that the critical amine group of RQ is not added in a late step of RQ synthesis but instead is present from the outset.

A key initial step in UQ synthesis is the addition by COQ-2 of a polyprenyl tail to a *p*-hydroxybenzoate ring (PHB — also often called 4-hydroxybenzoate (4-HB)) (Momose and Rudney, 1972; Trumpower et al., 1974). PHB has no amine group — however, *S.cerevisiae* COQ2 is known to be able to use a variety of similar compounds as substrates for prenylation such as para-aminobenzoic acid and vanillic acid (reviewed in Pierrel, 2017). Given the potential substrate flexibility of COQ-2, we hypothesized that RQ synthesis might start not with PHB but with a related molecule that contains an amine group already on the ring (Fig 3 for schematic). In particular, we noted that yeast COQ2 is tolerant of substituents at positions 5 and 6 of the PHB structure (reviewed in Pierrel, 2017) suggesting that this might be feasible enzymatically. We considered different candidate molecules as amine-containing ring structures that might act as precursors for RQ and focussed on anthranilate and 3-hydroxyanthranilate (3HA) as likely sources of the amine-containing ring in RQ. Anthranilate and 3HA are made from the amino acid tryptophan via the kynurenine pathway (Heidelberger and Gullberg, 1948; Kotake, 1936) and *kynu-1* encodes the *C.elegans* kynureninase that is required for the generation of anthranilate and 3HA(Babu, 1974; Bhat and Babu, 1980; van der Goot et al., 2012). We examined the quinones present in *kynu-1(e1003)* mutants that lack kynureninase and while UQ levels are normal in the *kynu-1(e1003)* mutant animals, there is no detectable RQ (Fig 3c). This suggests that the amine group on RQ derives from anthranilate or 3HA, or some closely related product of kynureninase.

To further confirm that the amine group on RQ ultimately derives from tryptophan we tested whether a tryptophan-derived aromatic amino group is being incorporated into RQ. We fed *C.elegans* 15N-labelled bacteria for 3 generations either in the presence or absence of 14N tryptophan. As shown in Fig 4a, the sole source of any 14N incorporated into RQ is the 14N tryptophan. As expected, the RQ detected in animals fed with 15N bacteria alone is approximately all 15N labelled (Fig 4b). However, if 14N tryptophan was added, ~50% of the RQ observed was 14N RQ — the sole source for this 14N was the added tryptophan. Taken together, our data show that RQ does not derive from UQ, and that anthranilate, 3HA, or a related molecule deriving from tryptophan via the kynurenine pathway is the source of the amine group on the quinone ring of RQ. We propose that the pathway of RQ and UQ synthesis are largely the same — the key difference is the presence or absence of the amine group on the initial aromatic ring substrate for COQ-2.

**Figure 4:**
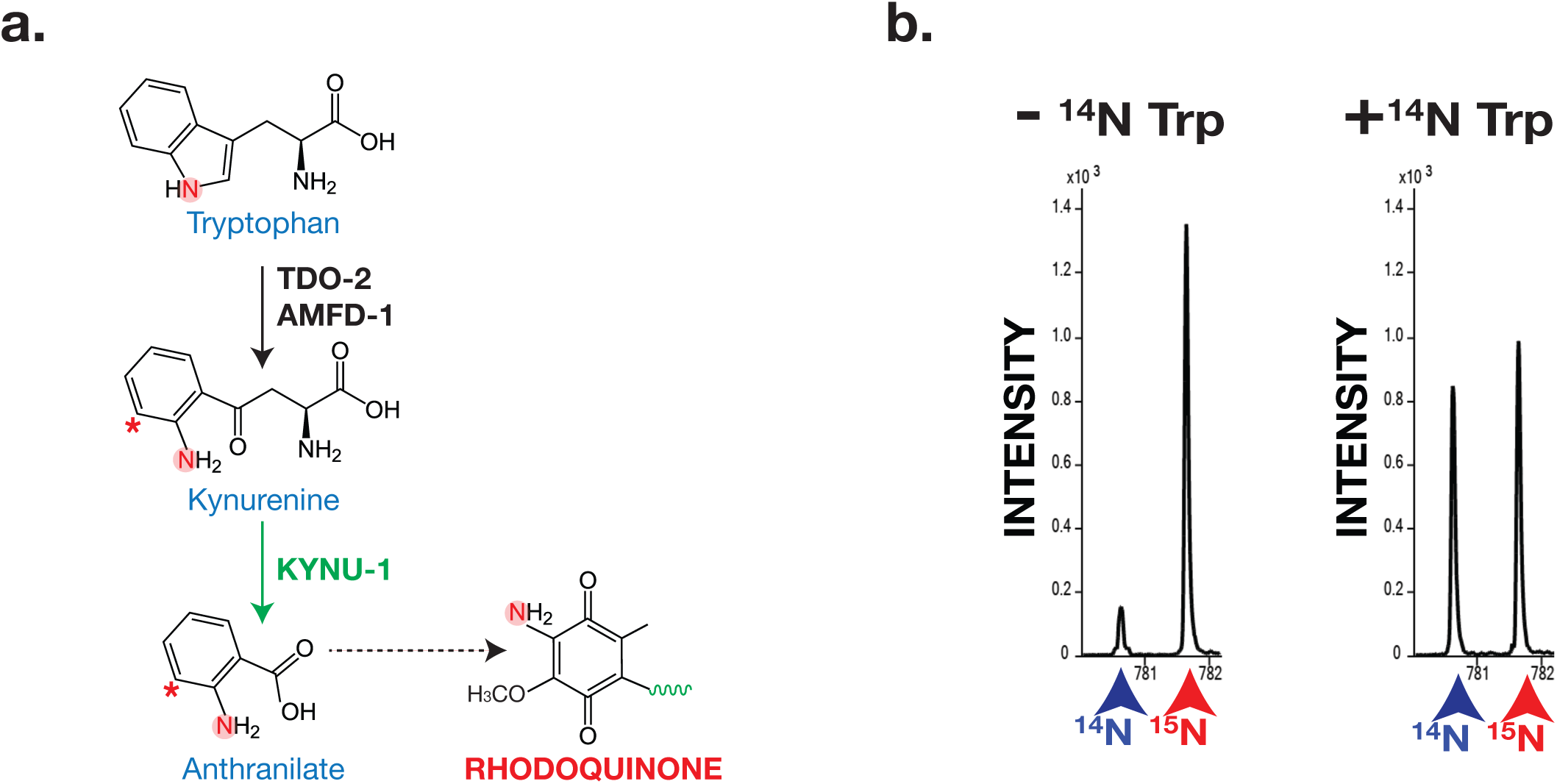
The critical amine group on RQ derives from Tryptophan. **a. Schematic of kynurenine pathway.** Anthranilate and 3-hydroxyanthranilate (3HA) derive from Tryptophan via the kynurenine pathway and this requires KYNU-1 activity. The nitrogen atom that becomes part of the key amine group on RQ is highlighted. **b. Analysis of RQ in animals fed either only 15N substrates or 15N substrates along with 14N Tryptophan.** Wild-type worms were fed with 15N-labelled bacteria for 3 generations either in the absence or in the presence of 14N Tryptophan. Almost all RQ is 15N labelled when worms were only eating 15N bacteria. However when 14N Tryptophan was also present, almost half the RQ is 14N RQ, indicating that the amine group of RQ must derive from Tryptophan.

### RQ is required for long-term survival of *C.elegans* in anaerobic conditions

As shown in Fig 2, *C.elegans* shows similar changes in metabolism when it is treated with KCN as STHs undergo when they adapt to the hypoxic environment of their host. For example, the classic hallmark of this RQ-dependent anaerobic metabolism is the generation of succinate by the action of Complex II as a fumarate reductase and *C.elegans* shows high levels of succinate when treated with KCN (Fig 3b). To confirm that this generation of succinate is indeed RQ-dependent in *C.elegans*, we examined whether RQ-deficient *kynu-1(e1003)* mutant animals could generate succinate when exposed to KCN. While wild-type worms generate high levels of succinate when Complex IV is inhibited with KCN, RQ-deficient *kynu-1(e1003)* mutant animals do not (Fig 5), confirming that the metabolic shift we see when we expose *C.elegans* to KCN is not simply similar to that of parasitic helminths in their hosts, it also requires RQ.

**Figure 5:**
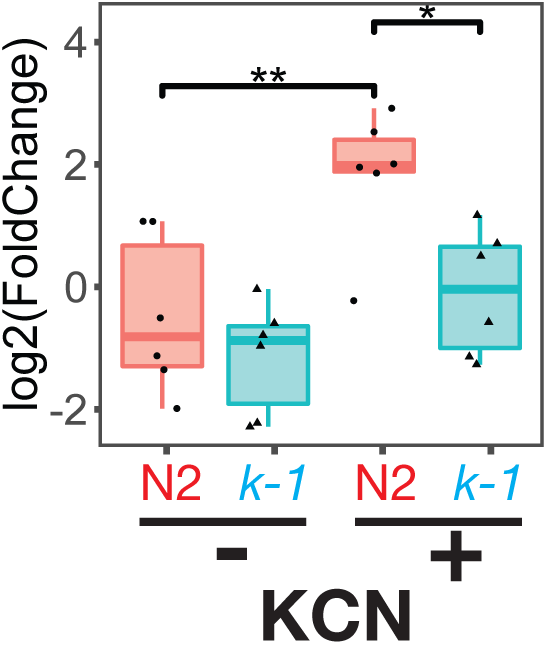
Buildup of succinate following Complex IV inhibition requires RQ. Wild-type (N2) or *kynu-1(e1003)* mutant animals were exposed to 200 µM KCN for 6 hrs and metabolites extracted and analysed by mass spectrometry. Data are from 5 independent biological repeats; box plots show median and the interquartile range as well as individual datapoints. Succinate levels increase markedly in N2 worms (red ‘N2’) following KCN treatment; there is no significant increase in *kynu-1(e1003)* mutant animals (cyan ‘*k-1*’) confirming that fumarate reductase activity of Complex II requires RQ.

We have thus established that treating *C.elegans* with KCN drives them into an alternative metabolic state where they use RQ to drive the same anaerobic metabolism used by STHs in their hosts. To be able to screen efficiently for drugs that affect RQ synthesis or RQ-dependent metabolism, however, we need a direct phenotypic readout for RQ-utilization rather than a molecular readout (such as succinate generation). When do *C.elegans* require RQ-dependent metabolism and what are the consequences if they have no RQ? Since RQ-dependent metabolism is being used in the presence of KCN we compared the sensitivity of wild-type worms and *kynu-1(e1003)* mutants to KCN and the ability of wild-type worms and *kynu-1(e1003)* mutants to survive in KCN for long periods. We found no significant differences in acute KCN sensitivity of wild-type worms and *kynu-1(e1003)* mutants — both slow their movement in the presence of KCN, stop moving completely by ~90 minutes (Fig 6a), and remain immobile from there on when maintained in KCN. However, there was a dramatic difference in their ability to survive extended periods in KCN. We exposed worms to KCN for different lengths of time and then removed animals from KCN and assayed their movement over the next 3hrs as they recover from KCN treatment. When wild-type worms are removed from KCN, they rapidly recover movement (Fig 6b) — they can do this even after 24 hours of KCN treatment (Fig 6c). However, *kynu-1(e1003)* mutants show greatly reduced ability to survive extended KCN treatment — they do not survive exposure to KCN for 12 hours or more (Fig 6c). RQ-dependent anaerobic metabolism thus allows *C.elegans* to survive extended periods where it cannot use oxygen as the terminal electron acceptor of the ETC.

**Figure 6:**
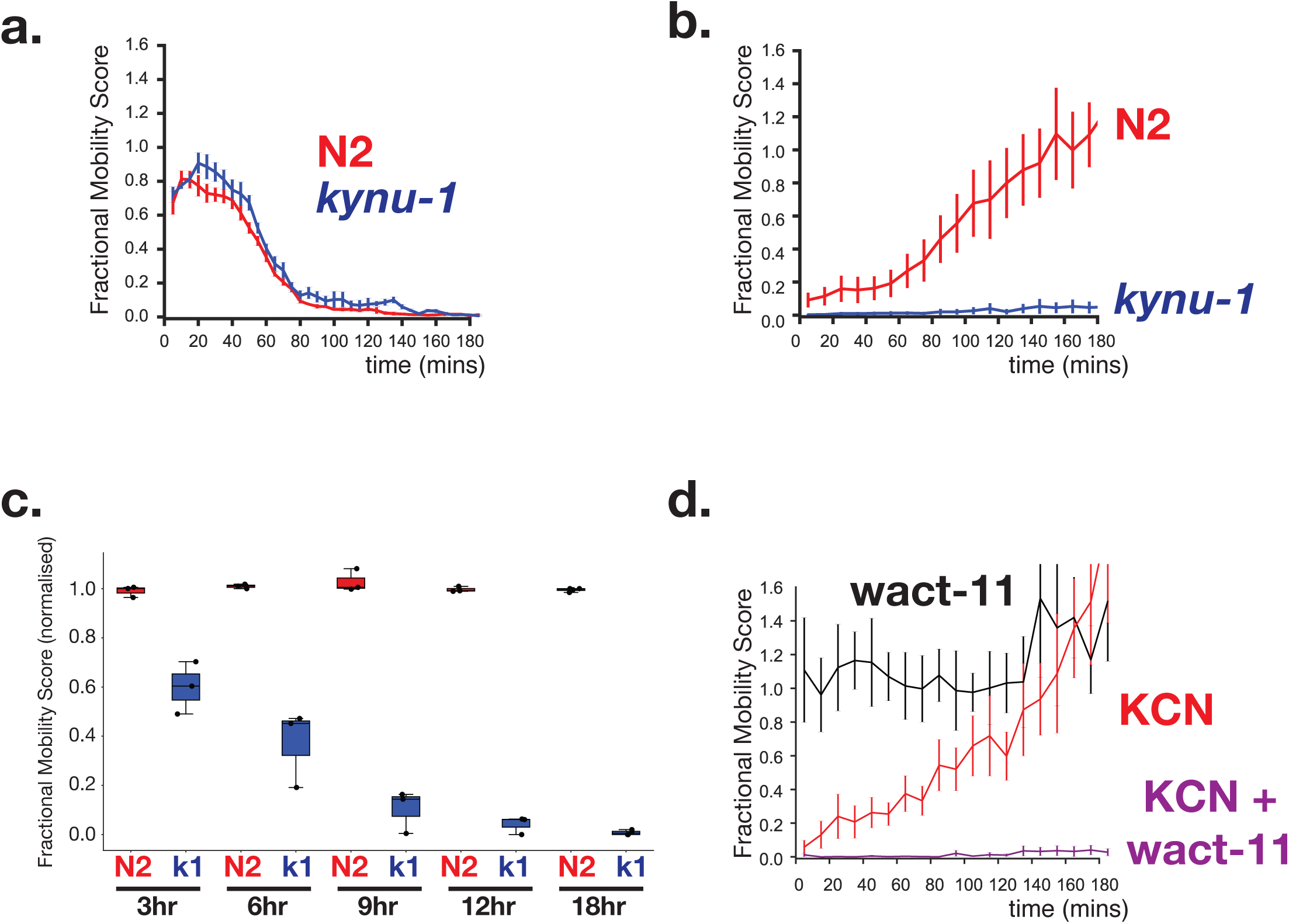
RQ-dependent metabolism is required for long-term survival in anaerobic conditions. **a. Loss of RQ does not affect acute sensitivity to KCN.** Wild-type (N2, red curve) and *kynu-1(e1003)* mutant L1 animals (*kynu-1*, blue curve) were exposed to 200 µM KCN and their movement measured over 3 hrs (see Methods). Both strains slow their movement and become immobile after ~90 mins. Curves show means of 3 biological replicates with 3 technical replicates in each; error bars are standard error. **b. RQ is required for survival following extended treatment with KCN.** Wild-type (N2, red curve) and *kynu-1(e1003)* mutant L1 animals (*kynu-1*, blue curve) were exposed to 200 µM KCN for 15 hrs. KCN was then diluted (see Methods) and worm movement was measured over a 3 hr time course. Curves show means of 3 biological replicates with 3 technical replicates in each; error bars are standard error. **c. Effect of different lengths of exposure to KCN on worm survival.** Wild-type (N2, red) and *kynu-1(e1003)* mutant L1 animals (*kynu-1*, blue) were exposed to 200 µM KCN for different lengths of time from 3 hr to 18 hr. KCN was then diluted (see Methods) and worm movement was measured after 3 hrs. Box plots show levels of movement after a 3 hr recovery period. Data are from 3 biological replicates with 3 technical replicates in each. **d. Ability to survive extended KCN exposure requires Complex II activity.** Wild-type L1 animals were treated with 200 µM alone, 10 µM of wact-11 (a helminth-specific Complex II inhibitor) alone, or a combination of KCN and wact-11 for 15 hrs. Drugs were then diluted 6x and worm movement was measured over a 3 hr timecourse. wact-11 treatment alone had no effect on survival over the experiment (black curve) and worms recovered completely from KCN treatment alone. However worms could not survive treatment with both KCN and wact-11. Curves show means of 3 biological replicates with 3 technical replicates in each; error bars are standard error.

This provides a simple assay for drugs that specifically affect RQ-dependent metabolism: drugs that block RQ synthesis or the activity of RQ-dependent pathways should abolish the ability of worms to survive >12hours in KCN. To test this, we used the compound wact-11 — this is a Complex II inhibitor that binds to the quinone-binding pocket of Complex II (Burns et al., 2015). wact-11 is highly related to the anthelmintic flutolanil (Burns et al., 2015) and is highly selective for helminth Complex II (Burns et al., 2015). Complex II is critical for RQ-dependent anaerobic metabolism where it acts as a fumarate reductase — inhibitors of Complex II might thus alter survival in KCN. That is what we observe: treatment with wact-11 prevents worms from surviving long-term KCN exposure (Figure 6d). Our assay will thus allow efficient screens for drugs that inhibit RQ synthesis or RQ utilization *in vivo* in a helminth under conditions where they require RQ, the first time this has been possible.

Finally, we took advantage of a set of mutant worm strains that are resistant to wact-11 treatment. Mutations that result in resistance to wact-11 treatment cluster in the quinone binding pocket of Complex II (Burns et al., 2015) and we reasoned that some of these might specifically disrupt the binding of RQ and thus affect the ability of worms to survive extended exposure to KCN. We tested a number of point mutants that affect wact-11 sensitivity (Fig 7a; data not shown) and found that most mutants appear similar to wild-type worms in their ability to survive long term KCN exposure. However, we found that the G71E mutation results in worms that are unable to survive extended KCN exposure — the G71E animals thus resemble *kynu-1(e1003)* mutants. We note that this mutation sits right above the modelled binding site for the rhodoquinone ring, whereas a neighbouring mutation that sits two turns of an alpha-helix further away (C78Y) has no effect. We thus suggest that the G71E mutation affects the ability of *C.elegans* to bind RQ into the quinone binding pocket of Complex II and thus to drive RQ dependent fumarate reduction as part of its RQ-dependent anaerobic metabolism.

**Figure 7:**
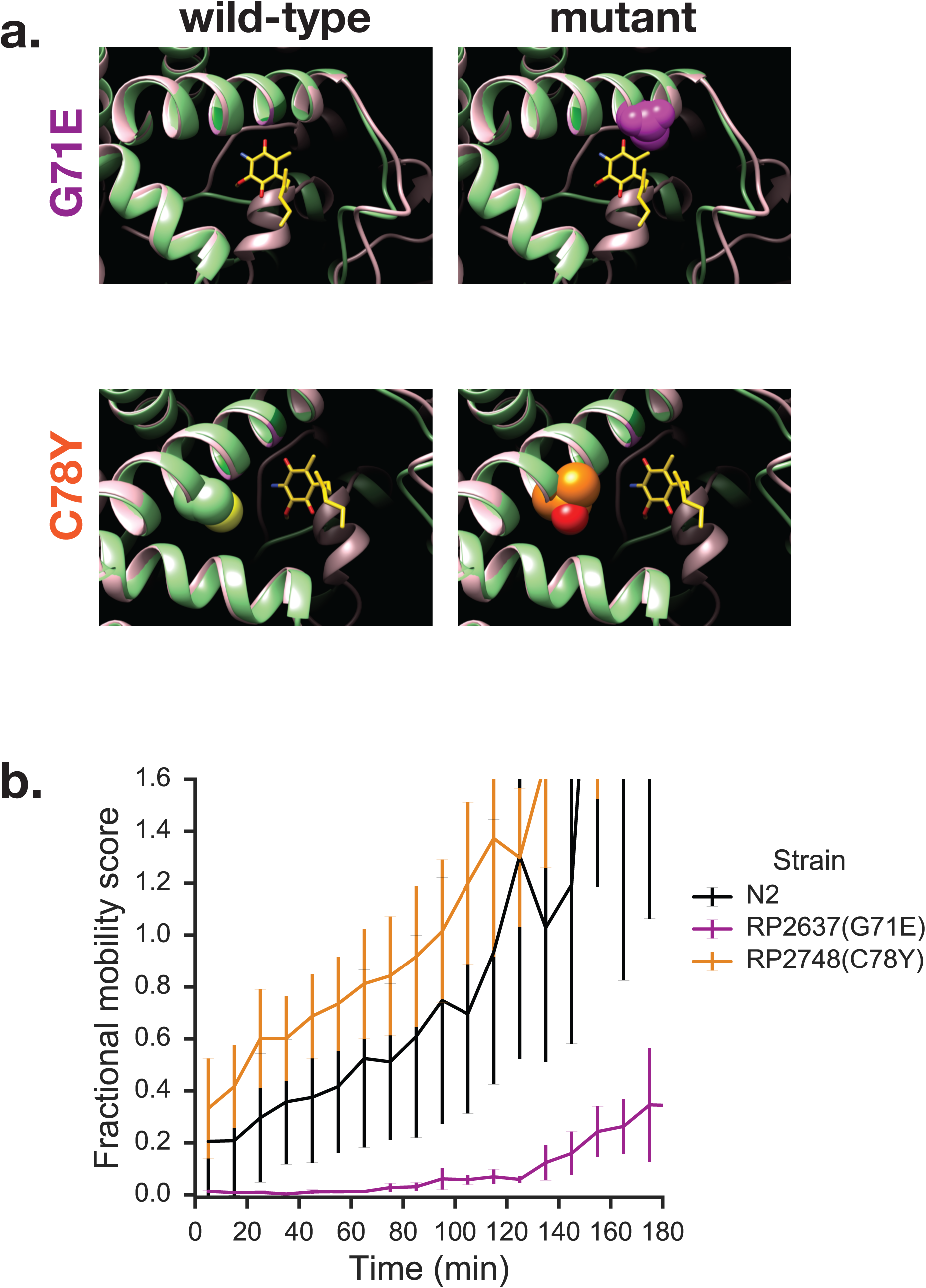
Mutations in quinone-binding pocket of Complex II affect ability to survive extended KCN treatment. **a. Positions of mutations that alter wact-11 sensitivity.** Structures show either the wild-type residues or mutations that alter wact-11 binding. The structures shown are *C.elegans* sequences (green) threaded onto the *Ascaris suum* crystal (pink). RQ is shown in yellow with the critical amine group in blue — note that this is RQ2 and not RQ9, hence the shortened prenyl tail. Note that G71E alters a residue that lies right over the quinone ring of RQ whereas C78Y is further from the ring. **b. Effect of quinone-pocket mutations on ability to survive extended KCN exposure.** L1 worms containing either wild-type, G71E or C78Y mutant *mev-1* sequences were exposed to 200 µM KCN for 15 hrs. KCN was then diluted 6x and the movement of worms measured across a 3 hr time course. Curves show means of 3 biological replicates with 3 technical replicates in each; error bars are standard error.

## Discussion

RQ was first identified over 50 years ago (Moore and Folkers, 1965). It is absolutely required for the survival of parasitic helminths in the hypoxic environment of the host gut where they can thrive for many months. The single amine group that differs between RQ and UQ is crucial for this — it allows RQ to carry electrons of the right electropotential to drive quinone-coupled dehydrogenases (QDHs) in reverse, acting as reductases (Fioravanti and Kim, 1988; Sato et al., 1972). In aerobic conditions, QDHs carry electrons from a diverse set of electron donors and transfer them onto UQ and hence into the ETC; under anaerobic conditions, RQ carries electrons to the QDHs which then reduce a diverse set of electron sinks, providing an exit point for electrons from the ETC. The single amine group on the quinone ring of RQ allows parasites to carry out this unusual anaerobic metabolism and thus it affects the lives of over a billion humans. Despite the importance of RQ for human health, its synthesis has been elusive and no anthelmintics have been identified that affect RQ synthesis. Here, we used *C.elegans* genetics to identify the RQ synthesis pathway and to establish a pipeline for screening for new compounds that alter the ability of worms to make and use RQ.

The critical question in RQ synthesis is where the critical amine group on the quinone ring comes from and how it is added. Previous studies suggested that RQ is synthesised using UQ as a precursor (Brajcich et al., 2010) and that the amine group is added at a late stage in RQ synthesis. Here, we show that this is not true, at least in *C.elegans*. We show that RQ synthesis does not require UQ as a precursor and, crucially, that the critical amine group on the quinone ring of RQ is not added at a late stage in the synthesis of RQ as has been previously proposed (Brajcich et al., 2010) but that it is present from the initial steps of RQ synthesis. For the first time since its discovery in the early 1960s, we now have a key insight into how RQ is made in helminths and this has several implications for the search for novel anthelmintics that might affect RQ synthesis.

First, we do not believe that there are separate dedicated pathways for UQ and for RQ synthesis in helminths. Instead, we suggest that UQ and RQ have a largely shared synthesis pathway. The key difference in RQ and UQ synthesis is the use of different initial substrates for COQ-2: if PHB is used, the product will be UQ, if anthranilate, 3HA, or a related product of tryptophan metabolism is used, the product will be RQ. This stands in clear contrast to the pathways used by a different set of organisms to make two different quinones, one for aerobic and one for anaerobic metabolism: facultative anaerobic bacteria including non-pathogenic *E.coli* as well as major human pathogens like *M.tuberculosis*. These bacteria make two quinones: UQ which is used as an electron carrier under aerobic conditions, and menaquinone (MK) (reviewed in Kwon and Meganathan, 2009), which acts as a carrier under anaerobic conditions and is in some sense analogous to RQ as it can carry electrons that can drive fumarate reduction (Cecchini et al., 1986). The synthesis pathways of UQ and MK are completely distinct and the genes involved are distinct (reviewed in Kwon and Meganathan, 2009). This separation of MK and UQ synthesis pathways has allowed the development of a number of promising compounds that act as inhibitors of the MK synthesis pathway (reviewed in Boersch et al., 2018). We suggest that there may be no analogous inhibitors for ‘the RQ synthesis pathway’ in helminths since there does not appear to be a dedicated RQ pathway analogous to the dedicated MK synthesis pathway.

Second, the pathway we identify for RQ synthesis suggests novel targets for anthelmintics. The finding that the kynurenine pathway is the source of the key precursors for RQ synthesis suggests naively that helminth-specific inhibitors of the kynureninase pathway might act as potent anthelmintics. However, the human gut is likely to be a source of anthranilate and 3HA from host metabolism or from the microbiome and inhibiting production of these molecules in the helminth itself might thus prove ineffective. A more likely target is COQ-2, the enzyme that prenylates the anthranilate or 3HA ring as the first step in RQ synthesis. Again, it is worth drawing an analogy between the synthesis pathways of UQ and MK in bacteria. The enzymes responsible for the prenylation steps of UQ and MK synthesis are entirely distinct and act at very different steps — UQ prenylation by *ubiA* (the *E.coli coq-2* orthologue) is a very early step, the MK prenylation by *menA* is a late step in MK synthesis (reviewed in Kwon and Meganathan, 2009). Inhibitors of *menA* (reviewed in Debnath et al., 2012; Kurosu et al., 2007) have no impact on *ubiA* or human COQ2 activity since menA and UbiA/COQ2 are completely different enzymes. Inhibitors that specifically block RQ synthesis in helminths while leaving host UQ synthesis intact will need to be more selective since they target orthologous enzymes, the human and helminth COQ2. However, since they have different substrate specificity, this may be possible and opens up a potential avenue for new anthelmintics.

Our study also raises key new questions. One of the most intriguing to us is why RQ synthesis is so rare amongst animals. To date, only three groups of animals are known to make RQ: molluscs, annelids, and helminths (Allen, 1973; Fioravanti and Kim, 1988; Klockiewicz et al., 2002; Sato and Ozawa, 1969; Takamiya et al., 2002; Van Hellemond et al., 1995). If RQ synthesis and UQ synthesis largely share a common pathway, why doesn’t every animal that makes UQ also make RQ? One possibility is that while much of the pathway for UQ and RQ synthesis is shared, RQ synthesis requires additional components that might only be present in RQ-synthesising species. We find some evidence that this may be the case.

The sole gene known to be required specifically for RQ synthesis and not for UQ synthesis is the *R.rubrum* gene *rquA* (Lonjers et al., 2012). This is a methyltransferase that is related to the quinone methyltransferases UbiG/COQ3 and UbiE/COQ5 that act in UQ synthesis (Lonjers et al., 2012; Stairs et al., 2018). Although RquA is predicted by homology to act as a quinone methylase (Lonjers et al., 2012), its exact role in RQ synthesis is still obscure and no clear orthologues in helminths or other RQ-producing animals have been identified to date. Whatever the function of RquA, it is clear that *R.rubrum* has three distinct quinone methyltransferases — two that are required for UQ synthesis and the third required specifically for RQ synthesis (Lonjers et al., 2012; Stairs et al., 2018). All animal genomes encode orthologues of UbiG/COQ3 and UbiE/COQ5 — these are the sole genes predicted to encode quinone methyltransferases in almost all animal genomes and there is no third related set like in *R.rubrum*. Helminths, molluscs, and annelids, the animal species known to make RQ, are different however — in addition to COQ3 and COQ5, they encode an additional set of related quinone methyltransferases (Fig 8a). In *C.elegans* this comprises four paralogues, *R08F11.4, R08E5.1, R08E5.3* and *K12D9.1*, and we find orthologues of these additional quinone methyltransferases in all parasitic helminths examined so far (examples in Fig 8a). Three of these additional quinone methyltransferase paralogues (*R08F11.4, R08E5.1* and *R08E5.3*) are highly upregulated in *C.elegans* mutants that are homozygous for a loss of function mutation in *gas-1* which encodes a subunit of Complex I in the ETC — *R08E5.3* is in the top 50 most upregulated genes, and all three are in the top 4% of upregulated genes (Falk et al., 2008). This suggests that their function may be required when aerobic respiration is compromised, as would be expected if they play a role in RQ synthesis. This correlation between expression of these additional quinone methyltransferases and RQ-dependent metabolism also appears to hold in parasites. For example, in *Ascaris*, RQ-dependent metabolism is essential for the survival of the adult worm and this is largely occurring in the muscle cells (Saz and Lescure, 1969). We identified an *Ascaris* orthologue of this additional class of quinone methyltransferases, *AgB01_g209*, that shows low expression in most developmental stages, but is strongly upregulated specifically in adult muscle cells (Fig 8b) (Jex et al., 2011). Thus while most animals only encode COQ3 and COQ5, RQ-synthesizing animal species encode an additional set of quinone methyltransferases whose expression is strongly upregulated under conditions requiring anaerobic respiration. This suggests that there may be similarity in RQ synthesis in *R.rubrum* and animals and that there may be at least some genes with functions specific to RQ synthesis. It will be interesting to examine whether these additional quinone methyltransferases participate in RQ synthesis in helminths and thus whether they are functionally analogous to *rquA* despite having only weak sequence similarity.

**Figure 8:**
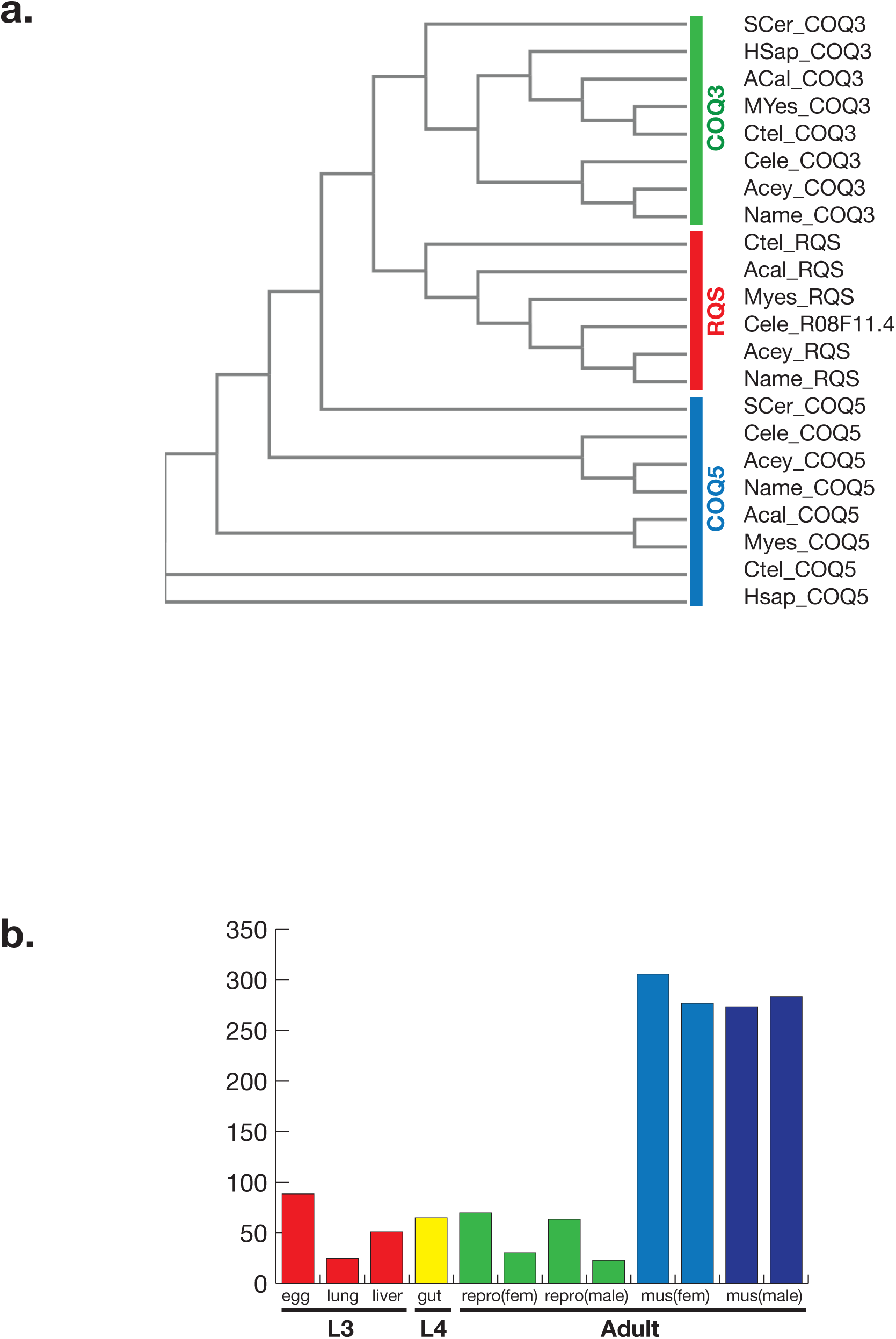
RQ synthesising species contain an additional class of quinone methyltransferases. **a. Eukaryotic quinone methyltransferases form three distinct classes.** We used BLAST to identify orthologues of UbiG/COQ3, UbiE/COQ5, and a novel additional class of quinone methyltransferases in *S. cerevisiae* (Scer)*, H. sapiens* (Hsap), the molluscs *A. californica* (Acal) and *M. yessoensis* (Myes), the annelid *C. telata* (Ctel) and the helminths *C. elegans, A. ceylanicum, N. americanus* (Cel, Acey, Name). Sequences were clustered using ClustalOmega (Sievers et al., 2011) and form three distinct clades: UbiG/COQ3, UbiE/COQ5, and the novel clade which we call RQS for RhodoQuinone Synthesis. **b. Developmental expression of *AgB01_g209*, an *Ascaris suum* orthologue of the novel quinone methyltransferases.** (Jex et al., 2011) Expression levels (in TPM) are shown for L3 stages (grown from isolated eggs *in vitro* (egg) or isolated from pig lung or liver), L4 stage larvae isolated from gut, and reproductive tissue or muscle tissue isolated from either male or female adult worms. Two adults are shown for each sample.

Finally, we note that several steps in the kynurenine pathway and in the ubiquinone synthesis pathway are catalysed by either monooxygenases or dioxygenases that require oxygen. These include TDO-2 (Hayaishi et al., 1957) and KMO-1 (Detmer and Massey, 1985; Entsch et al., 1976) in the kynurenine pathway and COQ-6 (Ozeir et al., 2015) and CLK-1 (Marbois and Clarke, 1996) in the UQ synthesis pathway. RQ synthesis thus appears to require the availability of oxygen for these enzymes, an unexpected result since RQ is preferentially required in anaerobic conditions and is the predominant quinone in helminths living under anaerobic conditions. How might these oxygen-requiring steps be carried out for RQ synthesis? It is possible that the helminth enzymes have evolved so that they can still operate under low oxygen conditions. Other oxygen-using proteins have evolved extremely high oxygen affinity in helminths — for example *Ascaris* haem is octameric and binds oxygen with ~25,000 times greater affinity than human haem (Minning et al., 1999). Alternatively, these same enzymatic steps might be carried out by other enzymes in lower oxygen conditions. In *E.coli*, for example, UbiB and UbiF (reviewed in Kwon and Meganathan, 2009) carry out the same hydroxylation modifications to the quinone ring as COQ-6 and CLK-1 in aerobic conditions and mutation of either gene results in a lack of mature UQ in these bacteria. However, under anaerobic conditions, *ubiB* and *ubiF* mutants make normal levels of UQ suggesting that other enzymes carry out these reactions in low oxygen conditions (Alexander and Young, 1978). It is possible that there is an analogous set of enzymes that are required for RQ synthesis in low oxygen conditions — these would carry out similar reactions to COQ-6 and CLK-1 but without the requirement for oxygen.

There is thus still much to be discovered about the regulation and the precise pathway of RQ synthesis in helminths. The results presented here provide a firm starting point and the assay we describe for drugs that affect RQ-dependent metabolism may lead to the discovery and development of a new class of anthelmintic drugs. Since resistance to known classes of anthelmintics is widespread among livestock parasites like *H.contortus*, *C.oncophora* and *A.suum* and is rising in human populations (reviewed in Sangster et al., 2018), this will prove critical in the control and treatment of these major pathogens.

## Materials and Methods

### Worm Strains and Maintenance

In addition to the traditional laboratory strain N2, here we include work using strains *clk-1(qm30), kynu-1(e1003), sdhc-1(tr357)*, and *sdhc-1(tr423)*. The two *sdhc-1* strains were provided by Dr. Peter Roy and all other strains were provided by the *Caenorhabditis* Genetics Centre. All worms were maintained on NGM agar plates seeded with *E. coli* OP50 as described elsewhere (Stiernagle, 2006) and maintained at 20°C.

### RQ Tryptophan Anti-Labelling Experiment

*Escherichia coli* (MG1655) was grown overnight at 37°C in M9 media prepared using 1 g/L ^15^N ammonium chloride (Cambridge Isotopes) as the nitrogen source. Bacteria were heat killed at 65°C for 15 minutes. The heated culture was used to seed NGM agar plates. 500 µL of 50 mg/ml tryptophan in water was spread on each 10cm plate. Ten L4 nematodes were placed on each plate to lay eggs overnight at 20°C. Adult worms were removed following the egg laying period. After 5 days at 20°C the nematodes were collected and frozen at −80°C.

### Quinone Extraction

Nematode samples were thawed and lysed via sonication. Quinone extraction solvent containing a 2:1 ratio of chloroform and methanol (Thermo Fisher Optima LC-MS grade) respectively was added to the samples. The organic phase of the sample was collected and then dried using nitrogen gas. Samples were resuspended in a 60:40 acetonitrile and isopropanol solution prior to analysis using APCI LC-MS.

### Quinone LC-MS analysis

Quinones were analyzed by reverse phase chromatography on an Eclipse Plus C-18 RRHD column, 2.1 mM x 50 mm with 1.8 um packing operated in a thermostatted column compartment held at 70°C. Buffer A was 50% MeCN in water, Buffer B was 100% acetone with 0.01% formic acid. Starting conditions were 0.25 mL/min at 50% B. Gradient was 1-minute hold, followed by increase 100% B at 5 minutes, hold 100% B until 7 minutes, then return to 50% B at 7.1 minutes and hold until 10 minutes. Samples were introduced from a HTC pal by injection of 5 µL sample into a 2 µL loop. Wash 1 was acetonitrile and wash 2 was isopropanol. Samples were ionized using a Multimode ionization source (Agilent) operated in APCI mode, gas temp 350°C, vaporizer temp 350°C, drying gas 5 L/min, nebulizer 60 PSI, capillary voltage 4000 V, corona current 4 µA, skimmer voltage 70 V, octupole 1 RF 400 V. Samples were analyzed on a 6230 TOF, a 6545 Q-TOF, or a 6490 QQQ as indicated. Fragmentor voltage for TOF/QTOF analysis was 200 V. For QQQ analysis, ubiquinone 9 was monitored by MRM of 795.6/197.3 at CID of 52 V; rhodoquinone-9 was monitored at 780.6/192.1 at CID of 52 V.

### Image-based assays

All image-based experiments were conducted on L1 animals which were collected from mixed-stage plates and isolated using a 96 well 11 µM Multiscreen Nylon Mesh filter plate (Millipore: S5EJ008M04) as described previously (Spensley et al., 2018) to a final concentration of ~100 animals per well. They were then incubated in a final concentration of 200 µM potassium cyanide for varying amounts of time. There are two key assays: the acute and the recovery. The acute assay monitors worm movement immediately following exposure to KCN every 5 minutes for a total of 3 hours. The recovery assays involve a KCN incubation of 3, 6, 9, 12, or 18 hours after which the KCN is diluted 6-fold with M9 buffer. Immediately after dilution, worm movement is monitored every 10 minutes for 3 hours. In both assays, worm movement is quantified using an image-based system as previously described (Spensley et al., 2018). All data were normalized by the fractional mobility score of the M9-only control wells per strain per time point.

### Drug preparation and Assay assembly

Solutions of potassium cyanide (Sigma 60178-25G) were made fresh prior to each experiment in phosphate buffered saline (PBS) and then diluted to a 5 mM stock solution in M9 buffer. 2X working concentrations were then prepared with M9 and the KCN stock solution. Wact-11 (Chembridge ID 6222549) was kept frozen as a 100 mM stock in DMSO. Diluted wact-11 stocks were made in DMSO (BioShop DMS666) to a concentration of 3.75 mM and kept frozen until day of use. 10X working concentrations were made with wact-11 stock, M9 buffer, and DMSO. All experiments were prepared to contain 0.8% v/v DMSO to control for any confounding effects of drug solvent.

Assays were assembled in flat-bottomed polystyrene 96-well plates (Corning 3997) to a total volume of 100 µL and 40 µL for the acute and recovery assays, respectively. Apart from assays including wact-11 which constituted half KCN solution, 10% wact-11 solution, and 40% worms in buffer, all other assays were comprised of equal parts worms in buffer and KCN solution.

### Rotenone LC-MS

Worms were collected and isolated as described above. A final concentration of 7.5 L1s/10µL was treated with final concentrations of 12.5µL rotenone (Sigma R8875) in 0.8% DMSO and with 0.8% DMSO alone and with 100 µM KCN in M9 or with M9 alone, 20 mL altogether in 40 mL plastic containers (Blender Bottle 600271) and were on a shaker for 1hr at room temperature.

After 1hr, samples were poured over 0.2 µM Nylaflo nylon filter membranes (PALL 66604) over vacuum and once the supernatant had run through, the filter paper was placed in prepared 1.2 mL of 8:1:1 extraction solvent (MeOH, HPLC Grade (SA 34860); H_-2_O, HPLC Grade (Caledon 8801-7-40); CHCl_3_, HPLC Grade (SA 650498)) in a 1.5 mL microfuge tube on dry ice. Tubes were inverted 5 times then vortexed.

Samples were switched between −80°C and −20°C three times. Filters were then removed, and the tubes spun at 13,200 rpm at 4°C for 30 min. 1 mL of supernatant was transferred to a new tube and dried under dry N_2_ with <0.02% O_2_ at 5 PSI for 8 hrs. Each sample was reconstituted with 30 µL HPLC grade water as was prepared labelled yeast reference. Samples and reference were spun at 13,200 rpm at 4°C for 5 min. 10 µL sample and reference were placed in an LC-MS sample vial (Agilent 5190-2243, cap is Agilent 5185-5820) and were fast spun at 1,000 rpm at 4°C.

### *kynu-1* LC-MS

N2 and *kynu-1(e1003)* worms were washed and filtered as previously described. They were then placed in 1.5 mL microfuge tubes at 1.5 mL for a final concentration of 45 L1s, 300 µM KCN in M9 or M9 alone and were placed on a rotator for 6 hours at room temperature. After 6 hours, the samples were extracted and prepared as previously described.

### Succinate Analysis

Succinate was extracted from an IPRP method LC-MS run at an mzCenter of 117.0193 and a retention time of 690 s in the case of the *kynu-1* experiment and from an Acid method LC-MS run at an mzCenter of 117.0193 and a retention time of 141 s in the case of the rotenone experiment. Both sets of samples were normalized to a labelled yeast reference. In the case of the *kynu-1* experiment, they were further normalized to the median of all the extracted peaks for each sample. In both cases they were ultimately normalized to the mean unlabelled N2 sample treated with buffer or 0.8% DMSO. Plots were generated using {plotPeak}.

### Structural Analysis

Sequences of succinate dehydrogenase, subunit C from *R. rubrum* (UniProtKB: Q2RV42), *E.coli* (UniProtKB: P0A8Q0 and P69054), *C. gigas* (UniProtKB: K1R921), *C. elegans* (Wormbase: CE00598 and CE50785), *A. suum* (UniProtKB: F1LC27), *S. cerevisiae* (UniProtKB: P33421 and Q04487), *H. sapiens* (UniProtKB: O14521), *M. musculus* (UniProtKB: Q9CZB0), *S. scrofa* (UniProtKB: D0VWV4), *M. balamuthi* (UniProtKB: A0A0B5D2L2), *A. ceylanicum* (UniProtKB: A0A0D6M6A9), *W. bancrofti* (UniProtKB: J9F801), *L. loa* (UniProtKB: A0A1I7VAG2), *A. simplex* (UniProtKB: A0A0M3K8C7), *O. bimaculoides* (UniProtKB: A0A0L8IER1), *A. vulgaris* (UniProtKB: A0A0B7BS14), *C. clemensi* (UniProtKB: C1C048), *M. yessoensis* (UniProtKB: A0A210PU64) underwent ClustalOmega multiple alignment (Sievers et al., 2011) using the default settings on EMBL-EBI.

Mitochondrial rhodoquinol-fumarate reductase from *A. suum* bound with rhodoquinone-2 (PDB: 3VR8) was displayed on Chimera (Pettersen et al., 2004) and the *C. elegans* sequence was threaded by homology using Modeller (Sali and Blundell, 1993; Webb and Sali, 2016) and the MSA with 15 iterations.

### Alignment of Quinone Methyltransferases

Sequences of COQ3/UbiG, Coq5/UbiE and the RQS cluster were identified by BLAST (Altschul et al., 1990) in *S. cerevisiae, H. sapiens, A. californica, M. yessoensis, C. telata*, *C. elegans, A. ceylanicum* and *N. americanus*. The sequences were then aligned by ClustalOmega (Sievers et al., 2011) and the phylogenetic relationships are taken from ClustalOmega (Sievers et al., 2011).

## Acknowledgements

The research in this study was supported by CIHR grant 501584. We thank Prof. Peter Roy and multiple members of Fraser and Roy labs for insightful discussions, Prof. Brent Derry for intellectual stimulation, the *Caenorhabditis* Genetics Center for *C.elegans* strains, Olga Zaslaver and Angela Wong for their timeand help wuth mass spec analyses. Structural graphics and analyses performed with UCSF Chimera, developed by the Resource for Biocomputing, Visualization, and Informatics at the University of California, San Francisco, with support from NIH P41-GM103311

---

**Supp Figure 1:**
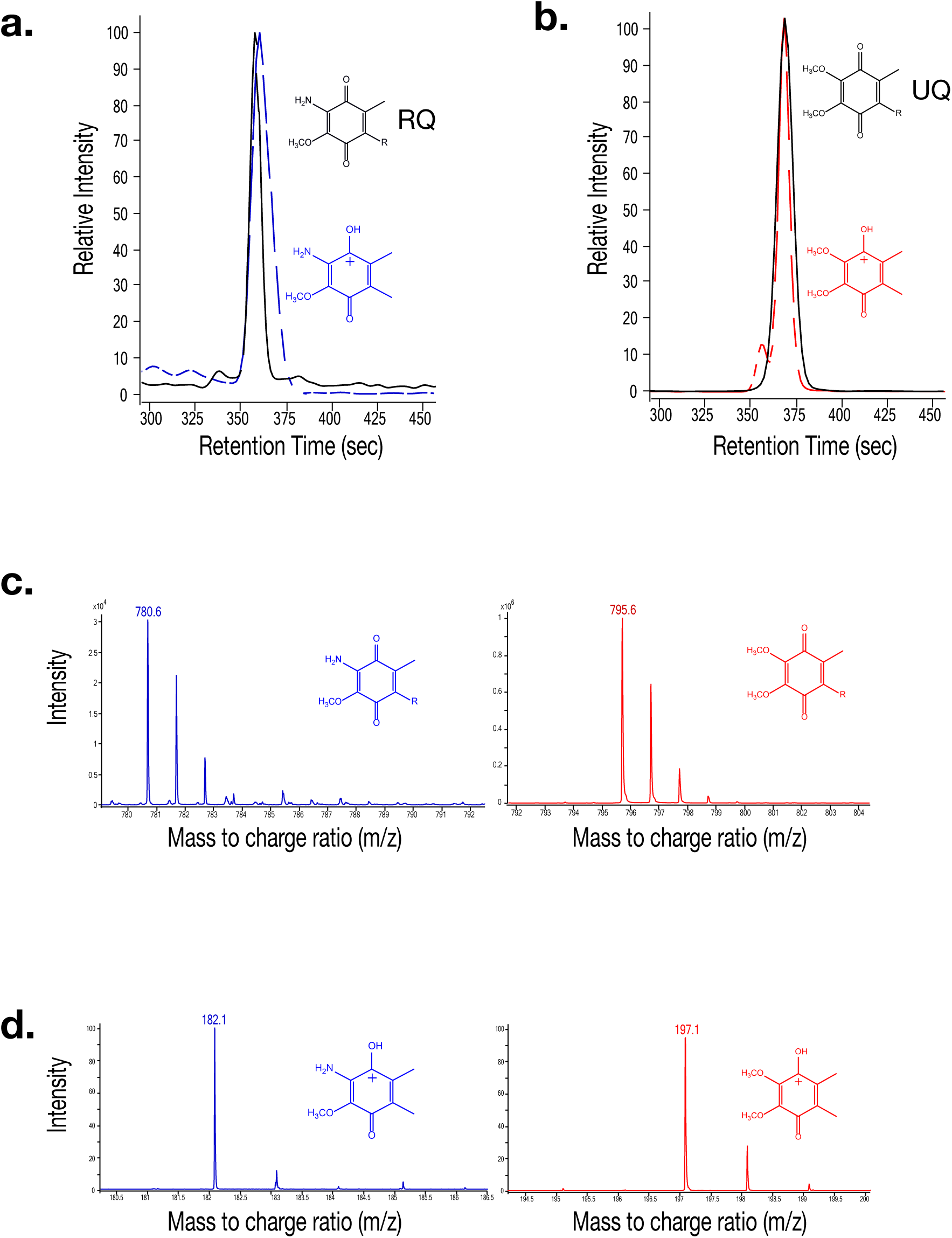
Mass spectra for Rhodoquinone (RQ) and Ubiquinone (UQ). **a.** Mass spectrometry (MS) extracted-ion chromatogram (EIC) of Rhodoquinone (RQ) (RT = 360 sec) shown by a solid black line. An MS/MS EIC of the RQ product ion is shown with a dashed blue line. **b.** MS EIC of Ubiquinone (UQ) (RT = 365 sec) shown by a solid black line. The dashed red line represents the MS/MS EIC of Q’s product ion. **c.** Mass spectra of RQ (m/z = 780.6) shown in blue. Mass spectra of UQ (m/z = 795.6) shown in red. **d.** Mass spectra of the RQ product ion (m/z = 182.1) shown in blue. Mass spectra of the Q product ion (m/z = 197.1) shown in red.

